## Supplementary material for "Motor response vigour and fixations reflect subjective preferences during intertemporal choice"

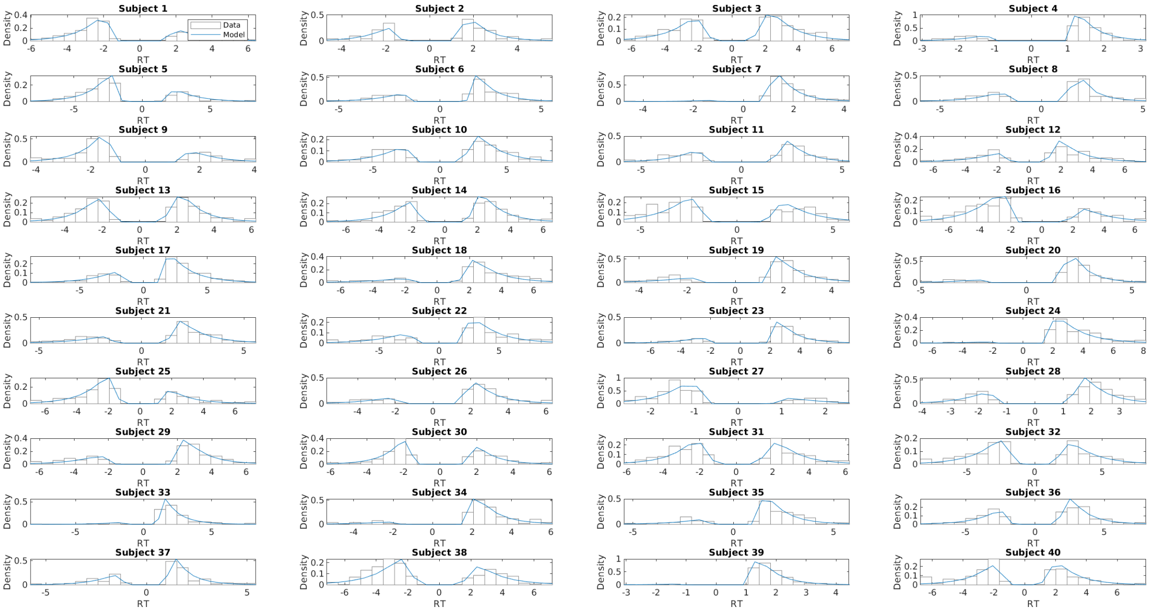


Figure S1: Posterior predictive response time distributions of the $DDM_{sig\text{-}shift}$ (using absolute values) for each participant, overlaid on the histograms of the observed RT distributions.

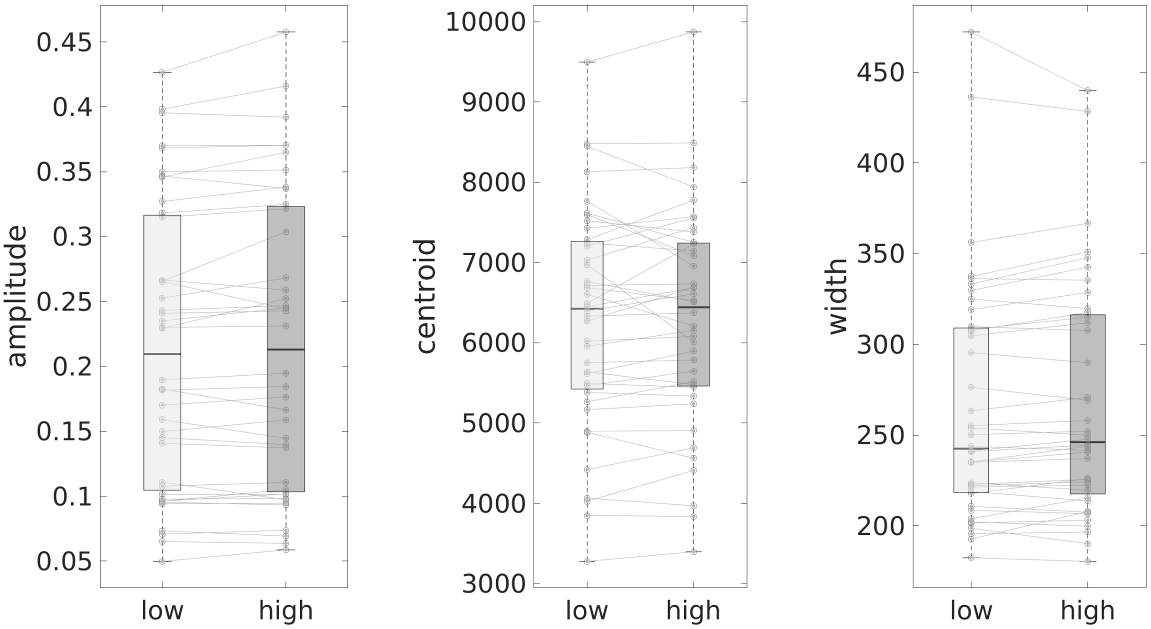


Figure S2: Parameters of the modelled grip response (mean values per participant and condition). The handgrip response was modelled with a Gaussian function with 1 term plus a constant (see Equation 10.

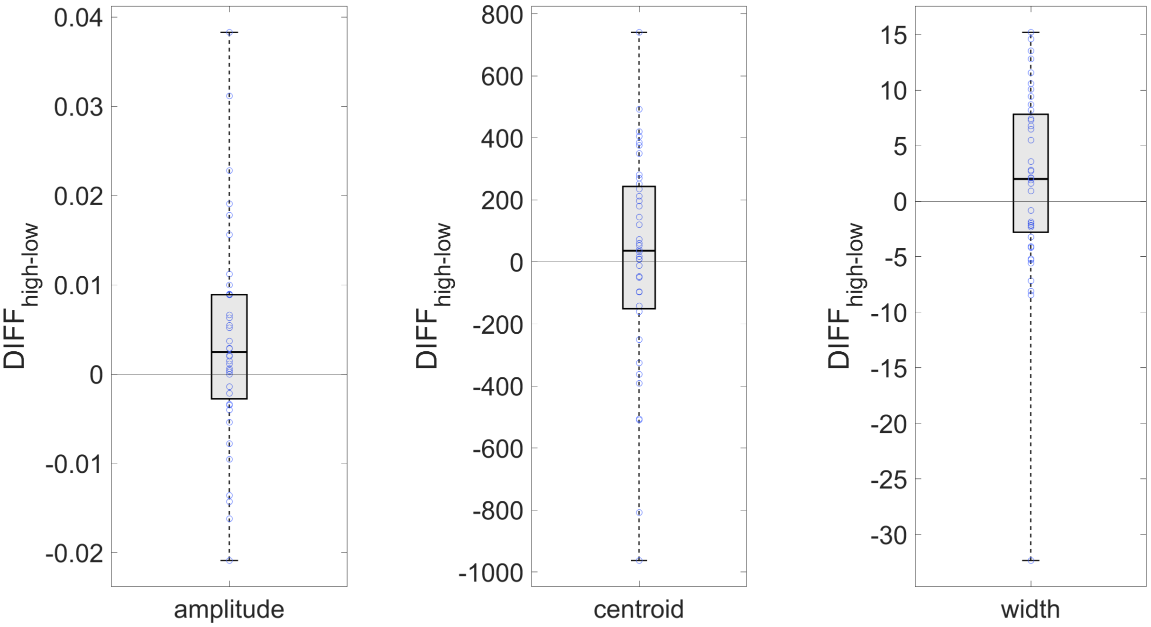


Figure S3: Within-subject differences of the parameters of the modelled grip response between the *low* and *high* condition. The handgrip response was modelled with a Gaussian function with 1 term plus a constant (see Equation 10).

### S1 Conflict based on trial-wise drift rate (DDM)

Response conflict was also operationalised based on the trial-wise drift rate calculated based on the estimated parameters of the highest-ranked DDM using normalised values ($DDM_{sig\text{-}shift}$). The posterior distributions of the group-level parameter means for the regression coefficients are depicted in Figure S4 (medians: $\alpha$ = 0.29 [intercept] $\beta_1$ = 0.03 [amplitude], $\beta_2$ = -0.02 [centroid], $\beta_3$ = -0.04 [width], $\beta_4$ = -0.05 [$N$ gaze shifts]). The Bayes factors provide only anectodal evidence that the regression coefficient for grip force amplitude is greater than zero vs. smaller than zero (BF for $\beta_1$: 1.29), or that the coefficients for centroid, width and number of gaze shifts are below zero rather than above zero (BF for $\beta_2$: 1.28, BF for $\beta_3$: 1.57, BF for $\beta_4$: 1.72). Since for all $\beta$s, the 95% HDI of the posterior distribution lie neither completely inside nor outside the ROPE, we remain undecided for these regression coefficients.

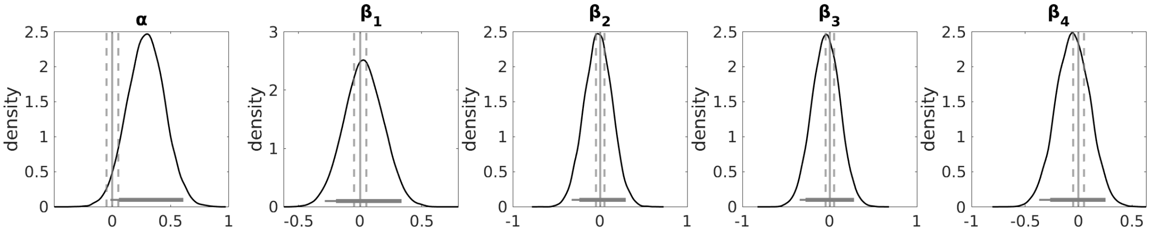


Figure S4: Hierarchical Bayesian regression results. Regression of the parameters of the Gaussian-modelled grip force response onto the trial-wise response conflict based on the drift rate (DDM). Posterior distributions of the group-level parameter means. $\alpha$: intercept, $\beta 1$: coefficient for amplitude, $\beta_2$: coefficient for centroid, $\beta_3$: coefficient for width, $\beta_4$: coefficient for fixation shift. Horizontal solid lines indicate the 85% and 95% highest density interval. Vertical solid lines indicate x = 0, and vertical dashed lines indicate the lower and upper bounds of the region of practical equivalence (ROPE).

### S2 Conflict based on subjective value differences (DDM using normalised values)

The medians of the group-level posterior distributions were as follows: $\alpha$ = 0.16 (intercept) $\beta_1$ = 0.01 (amplitude), $\beta_2$ = -0.02 (centroid), $\beta_3$ = 0.001 (width), $\beta_4$ = -0.004 ($N$ gaze shifts). The Bayes factors for all $\beta$ regression coefficients (for amplitude, centroid, width of the grip response and number of fixation shifts) provide only anectodal evidence for values greater than zero (BF for $\beta_1$: 1.07, BF for $\beta_2$: 0.78, BF for $\beta_3$: 1.02, BF for $\beta_4$: 0.89). We remain undecided for all $\beta$ coefficients, since the 95% HDI of the posterior distribution is neither completely inside nor outside the ROPE (see Figure S5).

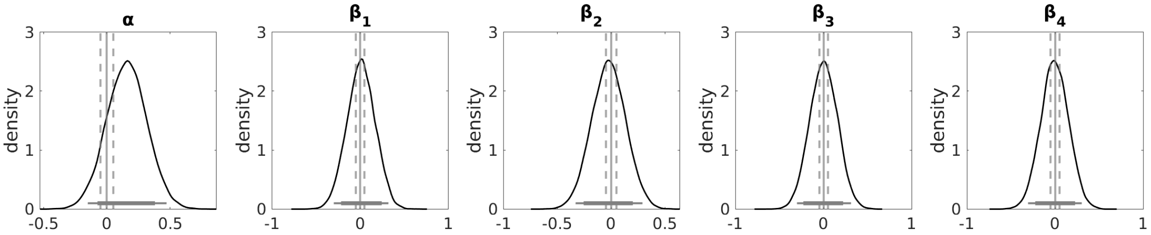


Figure S5: Hierarchical Bayesian regression results. Regression of the parameters of the Gaussian-modelled grip force response onto the value differences (DDM). Posterior distributions of the group-level parameter means. $\alpha$: intercept, $\beta1$: coefficient for grip force amplitude, $\beta_2$: coefficient for grip force centroid, $\beta_3$: coefficient for grip force width, $\beta_4$: coefficient for fixation shift. Horizontal solid lines indicate the 85% and 95% highest density interval. Vertical solid lines indicate x = 0, and vertical dashed lines indicate the lower and upper bounds of the region of practical equivalence (ROPE).
